# Characterizing microglial signaling dynamics during inflammation using single-cell mass cytometry

**DOI:** 10.1101/2024.07.27.605444

**Authors:** Sushanth Kumar, August D. Kahle, Austin B. Keeler, Eli R. Zunder, Christopher D. Deppmann

## Abstract

Microglia play a critical role in maintaining central nervous system (CNS) homeostasis and display remarkable plasticity in their response to inflammatory stimuli. However, the specific signaling profiles that microglia adopt during such challenges remain incompletely understood. Traditional transcriptomic approaches provide valuable insights, but fail to capture dynamic post-translational changes. In this study, we utilized time-resolved single-cell mass cytometry (CyTOF) to measure distinct signaling pathways activated in microglia upon exposure to bacterial and viral mimetics—lipopolysaccharide (LPS) and polyinosinic-polycytidylic acid (Poly(I:C)), respectively. Furthermore, we evaluated the immunomodulatory role of astrocytes on microglial signaling in mixed cultures. Microglia or mixed cultures derived from neonatal mice were treated with LPS or Poly(I:C) for 48 hrs. Cultures were stained with a panel of 33 metal-conjugated antibodies targeting signaling and identity markers. High-dimensional clustering analysis was used to identify emergent signaling modules. We found that LPS treatment led to more robust early activation of pp38, pERK, pRSK, and pCREB compared to Poly(I:C). Despite these differences, both LPS and Poly(I:C) upregulated the classical activation markers CD40 and CD86 at later time-points. Strikingly, the presence of astrocytes significantly blunted microglial responses to both stimuli, particularly dampening CD40 upregulation. Our studies demonstrate that single-cell mass cytometry effectively captures the dynamic signaling landscape of microglia under pro-inflammatory conditions. This approach may pave the way for targeted therapeutic investigations of various neuroinflammatory disorders. Moreover, our findings underscore the necessity of considering cellular context, such as astrocyte presence, in interpreting microglial behavior during inflammation.

**Main Points:** Time-resolved single cell mass cytometry delineates microglial signaling pathways following LPS or Poly(I:C) treatment. Astrocyte presence led to selective reduction of key microglial signaling nodes along with terminal inflammatory profiles.

## Introduction

### Microglial activation during pathology

Microglia, the resident immune cells of the central nervous system (CNS), play an indispensable role in maintaining tissue homeostasis. Under basal conditions, microglia exhibit a ramified morphology with elongated processes that facilitate continuous surveillance of the CNS environment. Upon activation—triggered by factors such as injury, infection, or neurodegeneration—these cells undergo a transformative change. The hallmark of this "activation" is a morphological shift where processes retract and thicken, adopting a bushier appearance. Notably, these gross morphological changes are accompanied by profound changes in microglial transcriptomes and proteomes that enable context-specific responses (1–3). Numerous studies have investigated transcriptional shifts in microglia during various pathophysiological conditions, from neurodevelopmental contexts to neurodegenerative diseases like Alzheimer’s (4–6). While these responses can be neuroprotective in certain situations, such as promoting tissue repair, they can also contribute to disease progression (7–10). Indeed, pharmacological microglial depletion has been shown to exacerbate viral encephalitis (7–8) but mitigate neurodegeneration in some models (9–10), underscoring the need for a nuanced understanding of microglial behavior during pathology.

### Molecular approaches to phenotype microglial activation

Traditionally, microglial activation has been categorized using the M1/M2 classification scheme, with M1 microglia being associated with pro-inflammatory and neurotoxic responses and M2 microglia with anti-inflammatory and tissue-reparative functions (1). However, this dichotomous framework fails to capture the full spectrum of microglial phenotypes. The advent of single-cell RNA sequencing (scRNA-seq) has revolutionized our understanding of microglial heterogeneity in both developmental and disease processes (4–6). Nevertheless, transcriptomics alone provides an incomplete picture, as mRNA levels often do not faithfully reflect protein abundance and post-translational modification (11–12). A comprehensive characterization of microglial states requires proteomic analysis, which can be achieved using single-cell mass cytometry, also known as CyTOF (Cytometry by Time-Of-Flight). This technique utilizes antibodies conjugated to heavy metal isotopes to enable simultaneous measurement of 40-50 affinity based measurement (e.g. proteins, phosphorylation events) (2,3,11,13). While previous CyTOF studies have examined microglial abundance and surface marker expression (2–3), an in-depth analysis of signaling dynamics is lacking.

### Modeling Inflammatory Challenges

To model microglial responses to pathogenic stimuli, lipopolysaccharide (LPS) and polyinosinic:polycytidylic acid (Poly(I:C)) are widely used to mimic bacterial and viral infections, respectively. LPS, a component of gram-negative bacterial cell walls, activates Toll-like receptor 4 (TLR4), while Poly(I:C), a synthetic double-stranded RNA analog, engages TLR3 (14). Although these compounds induce distinct responses at the morphological, transcriptomic, and cytokine levels (15–16), a comprehensive evaluation of the signaling pathways they activate in microglia remains lacking.

While previous studies have provided valuable insights into the transcriptional changes associated with microglial activation (4, 6, 15, 16), a comprehensive understanding of the dynamic signaling pathways that drive these responses remains elusive. Transcriptomic profiling offers a snapshot of cellular states but fails to capture the rapid post-translational modifications that propagate inflammatory signals (11, 12). While protein-based techniques such as western blotting and mass spectrometry provide a more direct read-out of cell function, these bulk approaches mask the heterogeneity of microglial responses, which can lead to oversimplification of their activation states. Single-cell mass cytometry (CyTOF) overcomes these limitations by enabling the simultaneous measurement of multiple signaling pathways and surface markers at single-cell resolution (2, 3, 11, 13). Recent CyTOF studies have revealed the existence of distinct microglial subsets in the context of aging and neurodegeneration (2, 3), highlighting the power of this approach to uncover novel biology. However, a detailed dissection of the signaling dynamics that underlie microglial responses to inflammatory stimuli is still lacking. Elucidating these pathways is crucial for identifying potential therapeutic targets and developing strategies to modulate microglial reactivity in neuroinflammatory disorders. To address this gap in knowledge, we employed a time-course CyTOF analysis to comprehensively map the signaling landscape of microglia following exposure to the bacterial endotoxin LPS and the viral mimetic Poly(I:C). Our results reveal stimulus-specific signaling trajectories, astrocyte-dependent immunomodulation, and a framework for rational drug design in neuroinflammation.

## Methods

### Mice

All experiments were carried out in compliance with the Association for Assessment of Laboratory Animal Care policies and approved by the University of Virginia Animal Care and Use Committee. Animals were housed on a 12 hr light/dark cycle. C57Bl6/J mice were purchased from Jackson (stock No. 000664) and were bred in-house to generate P0-P2 mice for culture experiments.

### Cell Culture

Briefly, mixed glial cultures were prepared from the cortices of newborn mice (P0–P2). Meninges were carefully removed in ice-cold DMEM/F12 (Thermo Fisher Scientific) and cortices dissociated via trituration. Dissociated cortices were spun at 600*g* for 5 minutes. Cells from 2 brains (*n* = 4 cortices) were plated in a T-75 flask coated with poly-d-lysine (Sigma) in 15 mL of DMEM/F12 (Thermo Fisher Scientific) with 10% FBS (Gibco), 1% penicillin and streptomycin (Thermo Fisher Scientific), sodium pyruvate (Thermo Fisher Scientific), and MEM Non-Essential Amino Acids (Thermo Fisher Scientific). On day in vitro 7 (DIV7) and DIV9, 5 mL of L-929 cell–conditioned medium (LCM) was added to promote microglial growth. To yield microglia-only cultures, mixed glial cultures were shaken on DIV12 - 14 at 180 rpm for 1 hour to dislodge microglia. Supernatants were centrifuged at 430g for 6 min and 250,000 microglia/well were plated in a 6 well-plate coated in poly-d-lysine. To yield microglia + astrocyte cultures, on DIV12 - 14 mixed glial cultures were trypsinized and plated at 400,000 cells/well in a PDK-coated 6 well plate. After two days, cultures were treated with either LPS (500 ng/mL) or Poly(I:C) (10 µg/mL) for varying amounts of time. Three independent time-courses per condition (microglia-only + LPS, microglia-only + Poly(I:C), microglia + astrocyte + LPS, microglia + astrocyte + Poly(I:C)) were generated for analyses.

### Mass cytometry Sample Dissociation

Following treatment, media was removed and then 1 mL of StemPro Accutase was added to the cells. Immediately upon Accutase addition cells were scraped off and added 1:1 to 4 % PFA. Cells were fixed for 10 min then spun at 500g for 5 minutes. Supernatant was discarded and cells were resuspended in 1 mL of 0.5 % BSA + 0.02 % sodium azide then stored in -80 C for later analysis.

### Mass Cytometry Sample Staining

Samples were stained and mass cytometry was performed as detailed in (11). Briefly, samples were barcoded by incubation with specific combinations of 1mM isothiocyanobenzyl EDTA-chelated palladium metals prior to being pooled into barcoded sets for staining. All time courses from both the microglia and microglia + astrocyte conditions were grouped together and stained with the same antibody master mix. Cells were first stained for surface epitopes by incubation with primary antibodies against extracellular proteins. Samples were next permeabilized with ice-cold 100% methanol prior to staining with primary antibodies against intracellular epitopes. To stabilize antibody staining, cells were then incubated with 1.6 % PFA alongside a DNA intercalator to aid in cell doublet and debris discrimination.

### Mass Cytometry

Cells were analyzed on a Helios CyTOF 2 system (Fluidigm). Prior to analysis, cells were resuspended in water (approximately 1 ml per 1L×L10^6^ cells) containing 1:20 EQ Four Element Calibration Beads (Fluidigm) and passed through a 40-μm nylon mesh filter. All samples were run simultaneously in one batch at a rate of 500 cells per second or less. Data were collected on a Helios CyTOF 2 using CyTOF Software version 6.7.1014.

### Normalization and Debarcoding

To control for variations in mass cytometer signal sensitivity across the run, raw .fcs files were normalized using EQ Four Element Calibration Beads (https://github.com/nolanlab/bead-normalization) (36). Normalized .fcs files from the run were then concatenated and debarcoded using software (https://github.com/zunderlab/single-cell-debarcoder) to deconvolute palladium metal expression on single cells according to a 6-choose-3 combinatorial system (37–38). A new parameter for barcode negativity (bc_neg) which is the sum of the three palladium measurements expected to be zero based on the cell barcode deconvolution assignment is then added to the .fcs files. Events with high bc_neg values likely contain two or more cells.

### Isolation of Single-Cell Events

To isolate single-cell events, normalized and debarcoded .fcs files were uploaded to Cytobank (https://community.cytobank.org) and clean-up gating was performed as described in Supplementary Figure 1. First, an additional debarcoding process was performed by removing events with a low barcode separation distance and/or high Mahalanobis distance. Singlets were then isolated by comparing the center of events with their lengths. Viable cells were then selected using a DNA intercalator dye. Next, events with a high cerium (Ce140) signal, indicating potential failure to remove a calibration bead during normalization, were removed. For high-dimensional analysis, these events were exported to .fcs files. For analysis of bulk microglial responses, an additional CD11b/CD45 gate was drawn to select for double-positive cells. To remove any potential residual astrocyte contaminants we selected for GFAP-negative cells. These events were then used to generate the data presented in Figure 1B-D, Supplementary Figure 1B-C, Figure 3B-C, Supplementary Figures 3 and 4.

**Figure 1.**
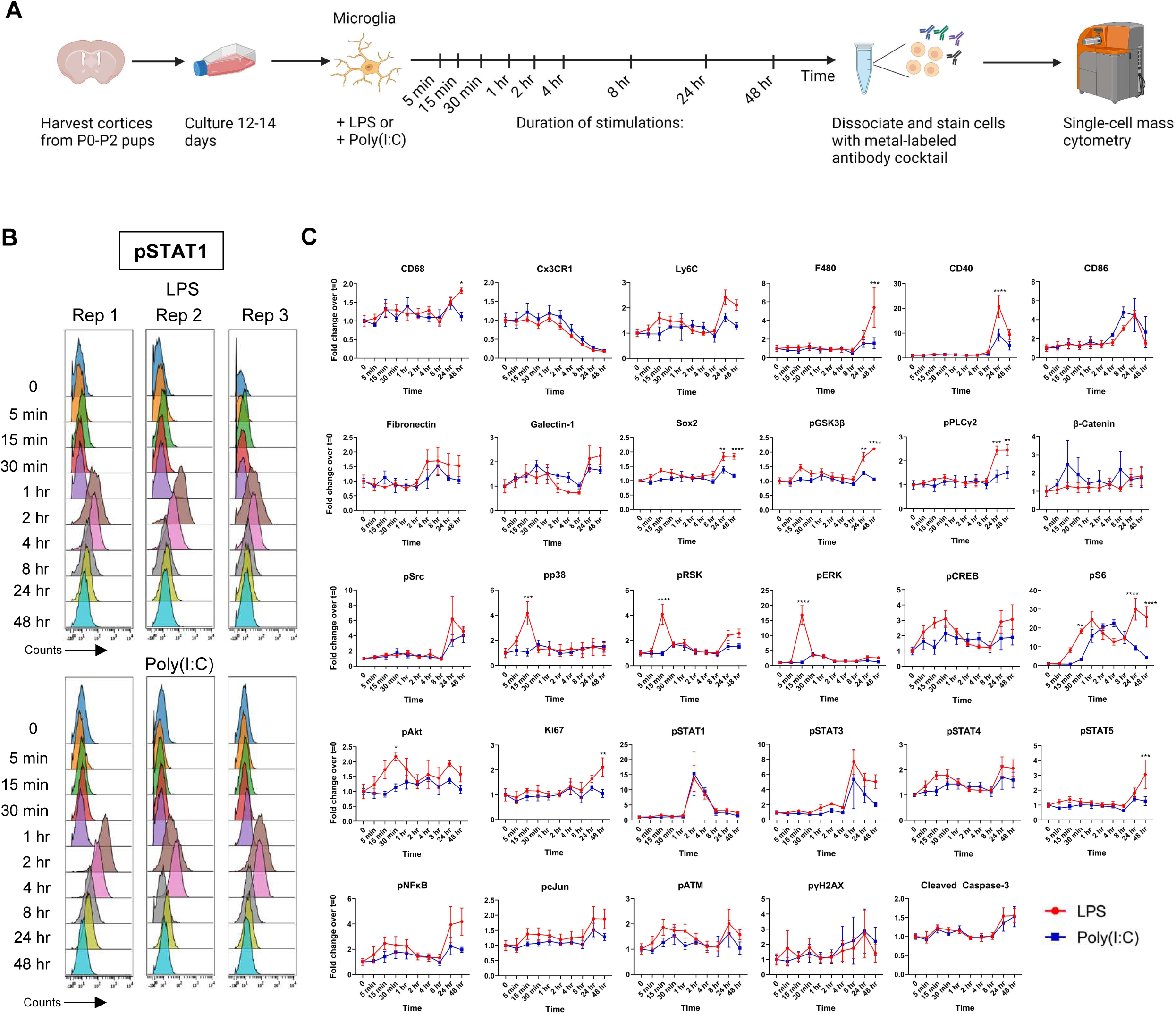
Microglia exhibit distinct signaling profiles when treated with LPS or Poly(I:C). Workflow for treatment paradigm and sample collection. Created using BioRender (A). pSTAT1 triplicates from LPS and Poly(I:C) time courses pulled from Supplemental Figure 1 to illustrate method reproducibility (B). Line-plots from LPS and Poly(I:C) treated microglial samples depicting signaling changes over time (C). Results are presented as fold-changes (normalized to respective vehicle intensity). Data were analyzed by two-way ANOVA followed by Šídák multiple comparisons test (C). Data are from at least three independent experiments and expressed as mean ± s.e.m. All data represent results taken from three independent cultures. Asterisks denote significance for LPS vs Poly(I:C) responses at a given time-point. *p < 0.05, **p < 0.01, ***p < 0.001, ****p < 0.0001.

**Figure 2.**
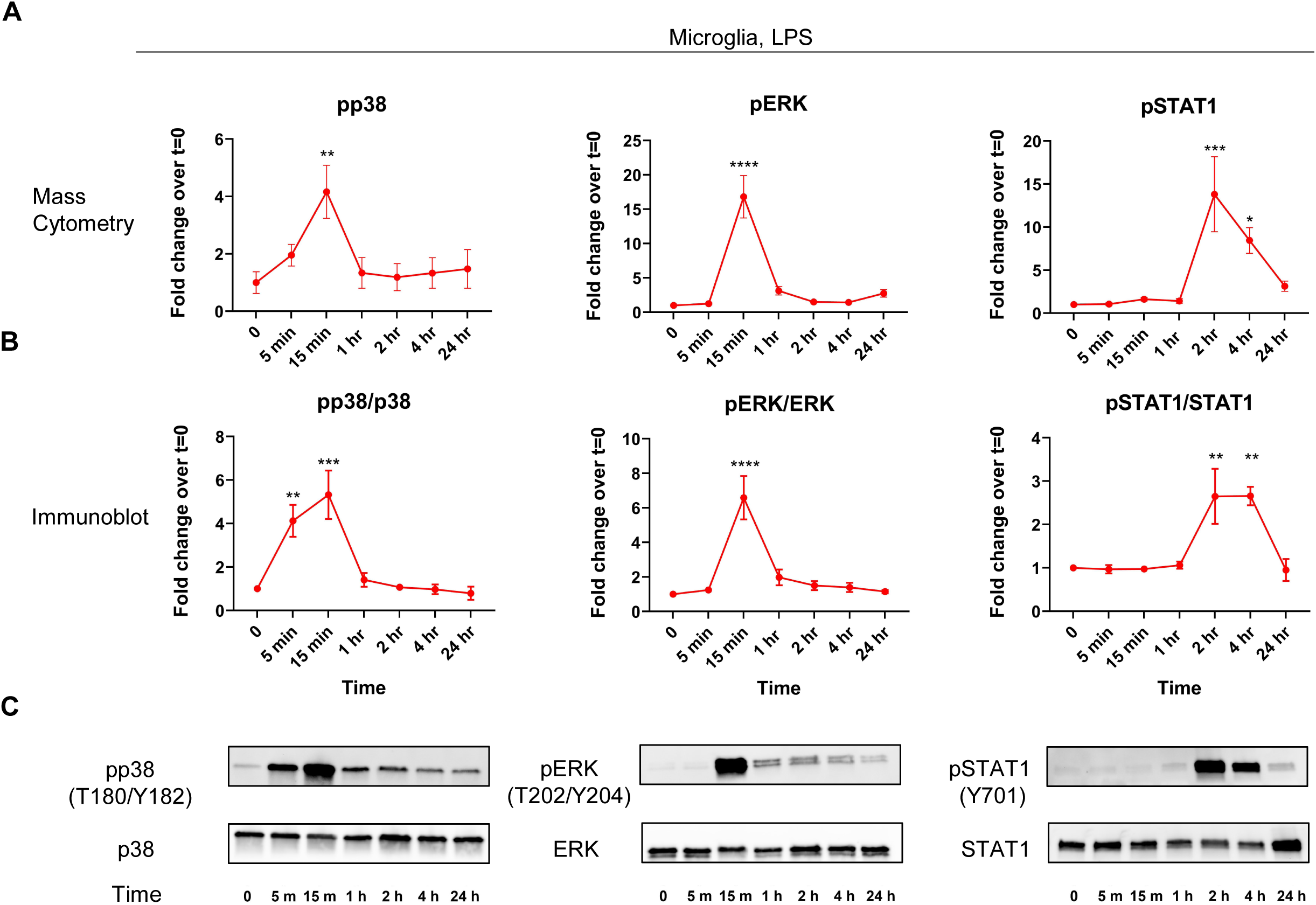
Western blot validation of mass cytometry signaling responses. Mass cytometry line-plots from LPS treated microglia for pp38, pERK, and pSTAT1 are replotted from Figure 1B for selected time-points (A). Western blotting quantifications for pp38, pERK, and pSTAT1 responses normalized to respective non-phosphorylated forms (B). Western blot images for data plotted in (B) (C). Data are from at least three independent experiments and expressed as mean ± s.e.m. All data represent results taken from three independent cultures. Asterisks denote significance for when fold-change is significantly changed from vehicle. (B) pp38/p38: 0 vs 15 min, **p = 0.0088. pERK/ERK: 0 vs 15 min, ****p < 0.0001. pSTAT1/STAT1: 0 vs 2 hr, ***p = 0.0008; 0 vs 4 hr, *p = 0.0432. (C) pp38/p38: 0 vs 5 min, **p = 0.0053; 0 vs 15 min, ***p = 0.0003. pERK/ERK: 0 vs 15 min, ****p < 0.0001. pSTAT1/STAT1: 0 vs 2 hr, **p = 0.0042; 0 vs 4 hr, **p = 0.0040.

**Figure 3.**
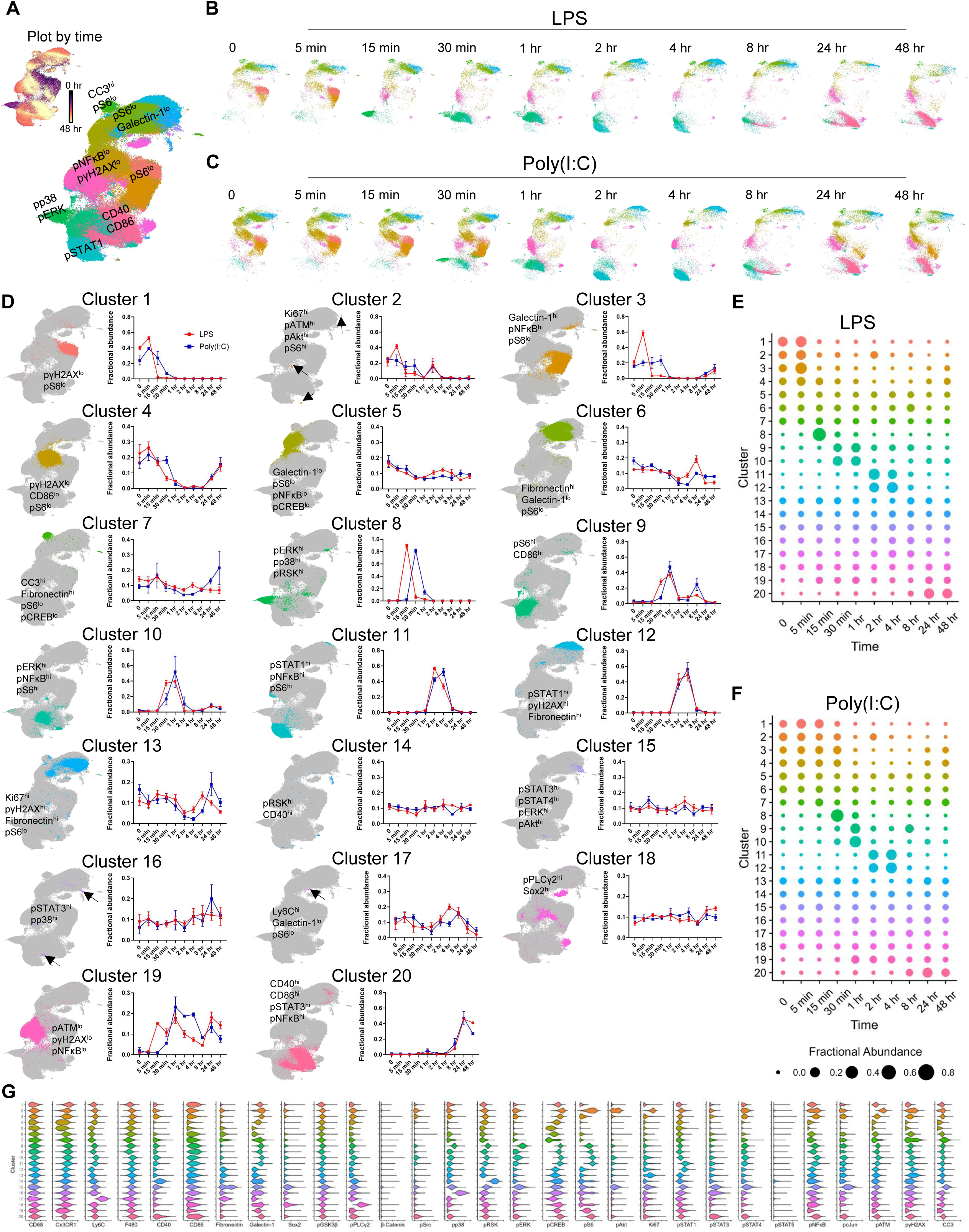
Characterizing LPS or Poly(I:C)-mediated microglial responses by clustering analysis. UMAP of all microglia (CD11b^+^CD45^+^ cells) with clusters annotated based on marker expression or colored based on time (Plot by time) (A). Same UMAPs as in (A) but separated based on LPS (B) or Poly(I:C) (C) treatment duration. Individual clusters from (A-C) are presented on grayed-out UMAP from (A) and annotated based on marker expression. Line plots showing changes in corresponding cluster abundance over time are presented to the right of the UMAP (D). Cow-plots displaying changes in cluster abundance over time for LPS (E) or Poly(I:C) (F) treatment. Cluster abundance is normalized with respect to time. As a result, clusters with greater abundance at early time-points are positioned in the upper left with clusters with greater abundance at later time-points positioned near the bottom right. Violin plots of protein expression from clusters in (D-F) (G). All data represent results taken from three independent cultures.

### Leiden Clustering

The Leiden community detection algorithm was used to partition cells into molecularly similar clusters (39). For the data presented in Figure 3 and Supplementary Figure 2A-B, events following cerium removal from microglia-only + LPS and microglia-only + Poly(I:C) (481,471 cells) were first clustered on identity markers (Olig2, CD11b, Fibronectin, GFAP, CD68, F480, CD45, Ly6C, Sox2, CD40, Galectin-1, Cx3CR1, and CD86) to identify microglial populations. We inspected violin plots for protein expression and excluded clusters with low CD11b, and/or low CD45, and/or high GFAP from further analysis. Olig2 expression was minimal across all clusters. A secondary round of clustering was then performed using all 33 markers in our panel except for CD11b, CD45, GFAP, and Olig2. A similar workflow was used for the data presented in Figure 3D-H, Supplementary Figure 2C-D, Supplementary Figure 5-6 which used events from microglia-only + LPS, microglia-only + Poly(I:C), microglia + astrocyte + LPS, microglia + astrocyte + Poly(I:C) exported .fcs files (758,033 cells).

### UMAP Data Visualization

Data was visualized by uniform-manifold and approximation (UMAP, https://github.com/lmcinnes/umap) with the following parameters: nearest neighbors = 15, metric = euclidean, local connectivity = 1, N components layout = 2, epochs = 1000 (40).

### Western Blotting

Following treatments, cells were washed with ice-cold 1x PBS then lysed in 1x RIPA buffer (25 mM Tris-HCl, pH 7.5, 150 mM NaCl, 1% NP-40, 0.5% sodium deoxycholate, 0.1% SDS) supplemented with with complete protease inhibitor (Roche) and PhosSTOP phosphatase inhibitor (Roche). Cells were kept on ice for 20 min and then centrifuged at 14,000g for 10 min at 4 C. 10 µg of lysate were boiled for 5 min in an equal volume of 2x laemmli buffer prior to loading onto a 4-15% Mini-PROTEAN TGX Precast Gel. Samples were then transferred onto PVDF membranes and blocked for 1 hr in 5 % milk (Fisher, catalog # NC9121673) in TBS + 0.05% Tween (TBST) at room temperature. Membranes were then probed overnight at 4 C with primary antibody in 5 % milk in TBST. Membranes were then washed in TBST then incubated for 1 hr at RT with the corresponding HRP conjugated secondary antibody in 5 % milk in TBST (1:5,000). Membranes were then washed again in TBS and signals developed using the SuperSignal™ West Femto Maximum Sensitivity Substrate (Thermo Fisher Scientific; according to the manufacturer’s protocol). Blots were stained first with anti-phospho antibodies then stripped using the Western BLot Stripping Buffer (Takara; according to the manufacturer’s protocol) and reprobed with the corresponding antibody towards the non-phosphorylated target. The following antibodies were used: anti-phospho-STAT1 (CST (9167S), 1:1000), anti-STAT1 (CST (9172T), 1:1000), anti-phospho-p38 MAPK (CST (4511T), 1:1000), anti-p38 MAPK (CST (8690T), 1:1000), anti-phospho-p44/42 MAPK (CST (9101S), 1:1000), anti-p44/42 MAPK (CST (9102S), 1:1000).

### Statistics

Data in figures represents mean ± SEM. Each n represents an independent biological sample. Analysis was performed on Graphpad Prism 10, applying a two-way ANOVA with a Šídák multiple comparisons test (Fig. 1C, Fig. 4C, Fig. 4H, Fig. 5B, Fig. S3A-B, Fig. S4A-B) or one-way ANOVA with a Dunnett’s multiple comparisons test (Fig. 2A-B).

**Figure 4.**
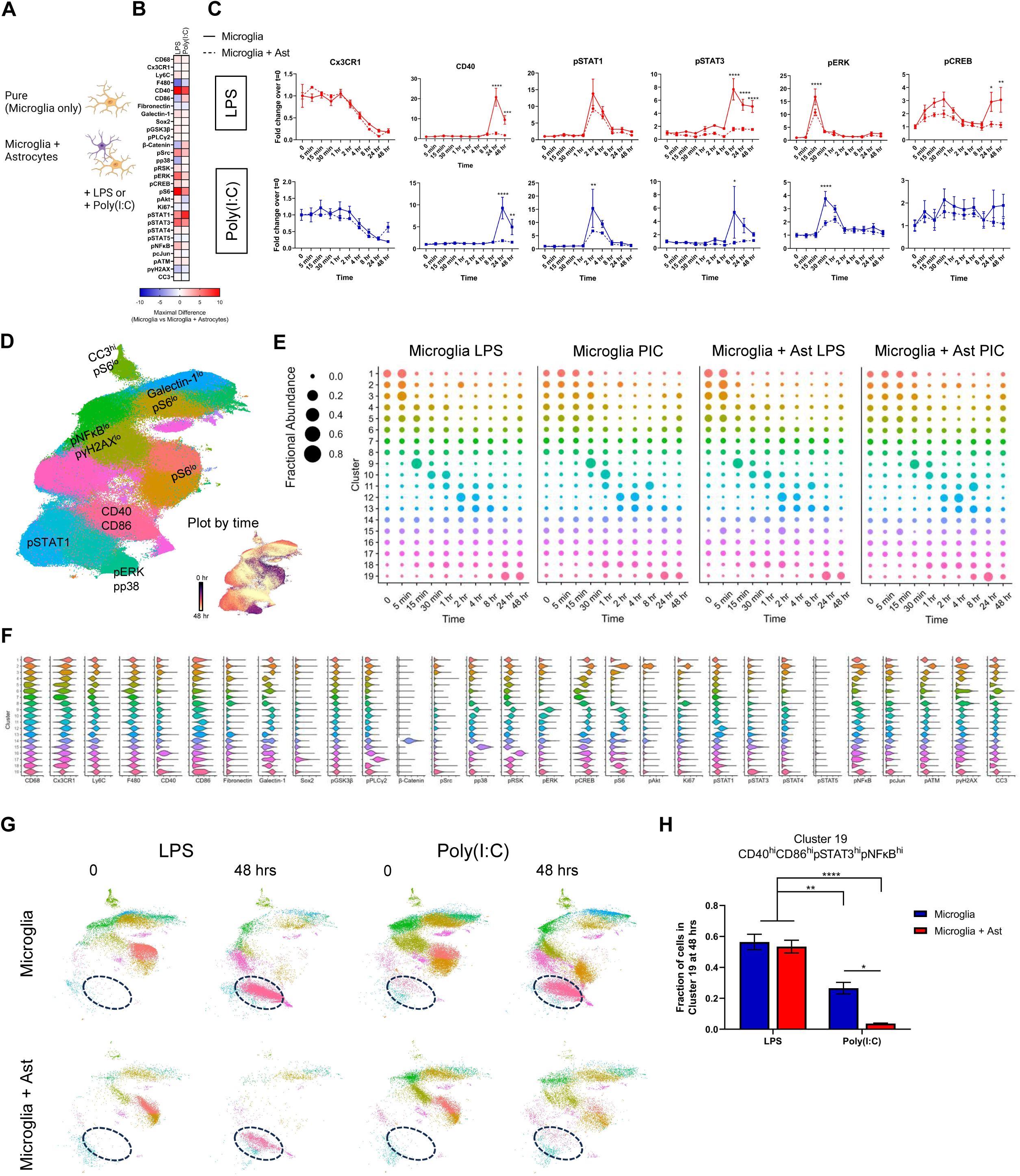
Effects of astrocytes on microglial signaling profiles. Microglial responses were compared either in the presence or absence of astrocytes (Microglia only vs Microglia + Astrocytes). Image created using BioRender (A). The difference in signaling marker fold-changes between microglia-only and microglia + astrocytes were calculated for each time-point. Heatmap displays the maximal difference with warmer colors indicating markers that were greater in the microglia-only setting and cooler colors indicating markers that were greater in the microglia+astrocyte setting (B). Select fold-change line-plots from LPS and Poly(I:C) treated microglia-only and microglia+astrocyte (C). UMAP of all microglia (CD11b^+^CD45^+^ cells) with clusters annotated based on marker expression or colored based on time (Plot by time). Cow-plots displaying changes in cluster abundance over time for respective treatment conditions (E). Violin plots of protein expression from clusters in (D) (F). UMAPs from 0 and 48 hr mark for microglia-only (LPS), microglia-only (Poly(I:C)), microglia + astrocyte (LPS), and microglia + astrocyte (Poly(I:C)) conditions with Cluster 19 circled (G). Fraction of cells in cluster 19 from (G) (H). Data were analyzed by two-way ANOVA followed by Šídák multiple comparisons test. Data are from at least three independent experiments and expressed as mean ± s.e.m. All data represent results taken from three independent cultures. (C) Asterisks denote significance for Microglia vs Microglia + Ast responses at a given time-point. *p < 0.05, **p < 0.01, ***p < 0.001, ****p < 0.0001.

**Figure 5.**
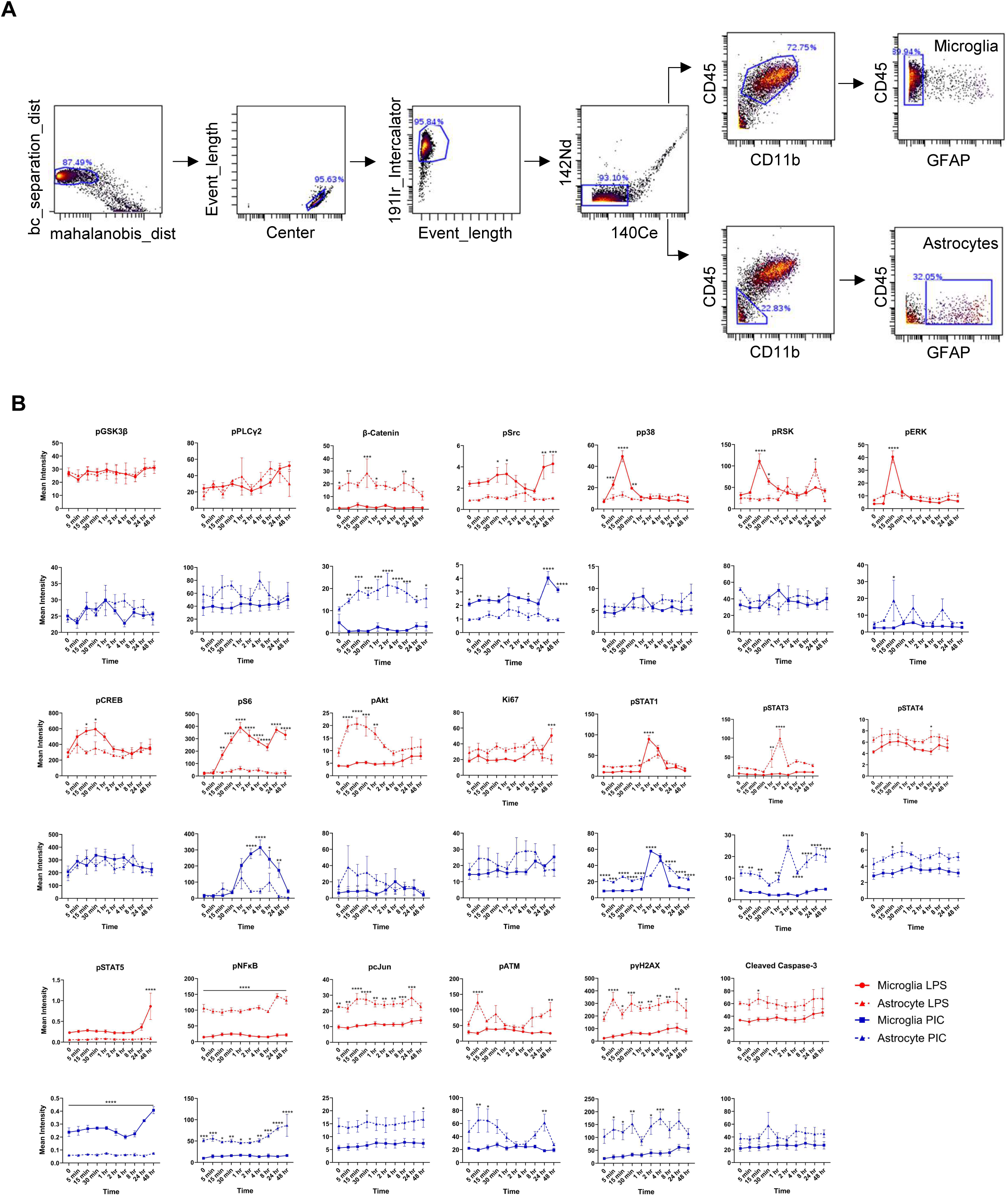
Astrocyte profiles following LPS or Poly(I:C) treatment. Cytobank gating strategy to isolate astrocytes (A). Mean marker intensity line-plots for astrocytes from LPS or Poly(I:C) time-courses. Microglial responses are also included for comparison (B). Data were analyzed by two-way ANOVA followed by Šídák multiple comparisons test (B). Data are from at least three independent experiments and expressed as mean ± s.e.m. All data represent results taken from three independent cultures. Asterisks denote significance for Microglia vs Astrocyte mean intensities at a given time-point. *p < 0.05, **p < 0.01, ***p < 0.001, ****p < 0.0001.

**Figure 6.**
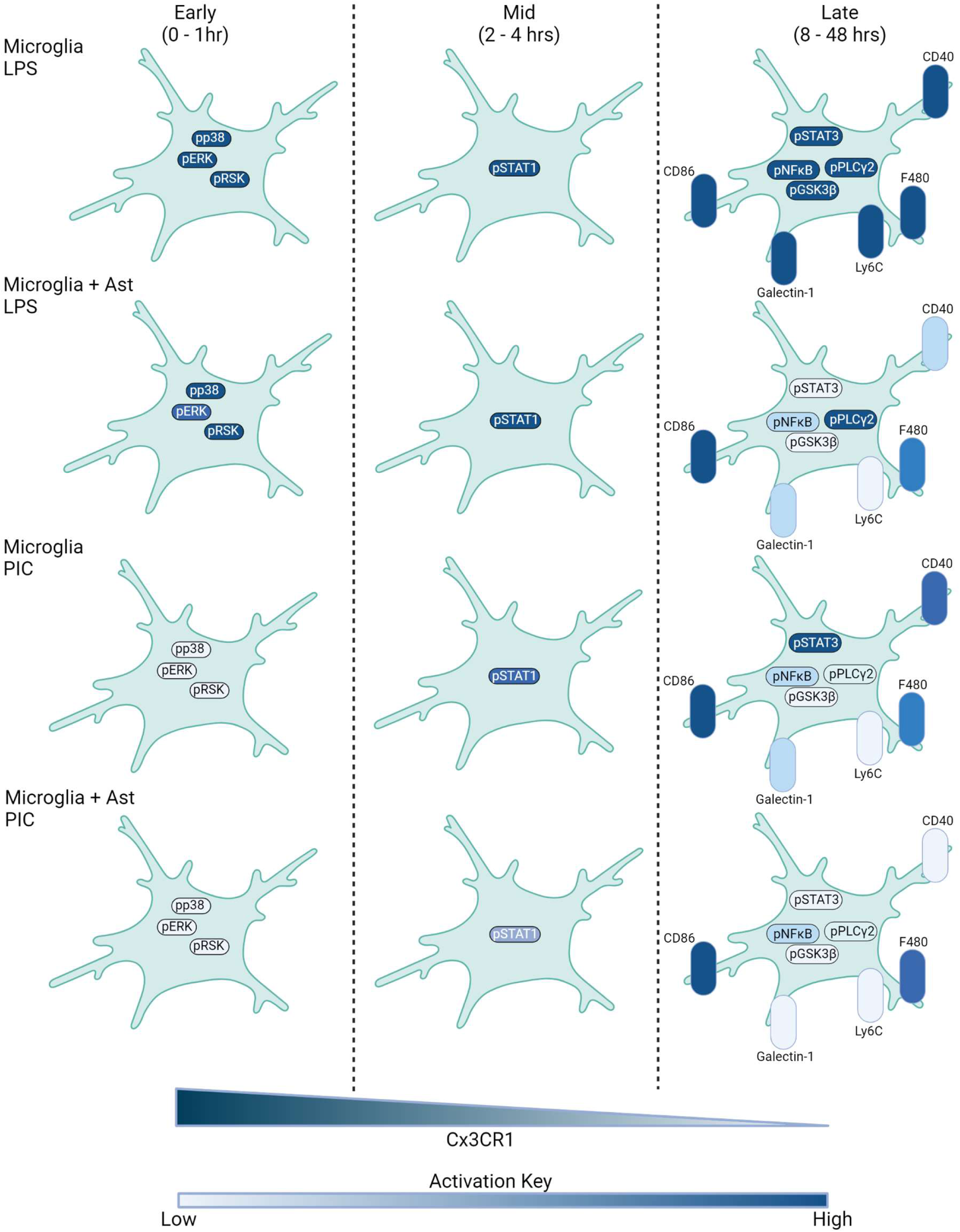
Summary of LPS and Poly(I:C) transitions along with astrocyte modulation. Major signaling changes during early (0 - 1 hr), mid (2 - 4 hrs), and late (8 - 48 hrs) stages of LPS and PIC treatment. Intensity of color depicts magnitude of marker change with darker colors indicating greater signal strength.

## Results

### Microglia exhibit distinct signaling profiles when treated with LPS or Poly(I:C)

To analyze the signaling dynamics of microglia when faced with pro-inflammatory insults, we cultured microglia from P0-P2 mice and challenged them with either the TLR4 agonist LPS or the TLR3 agonist Poly(I:C). Microglia were then harvested and stained with a metal-conjugated antibody panel comprising 33 markers targeting key signaling pathways and cell identity markers (21 signaling and 12 identity) (Figure 1A, Supplementary Table 1, Supplementary Figure 1A). At early time-points, CD11b^pos^CD45^pos^ (microglia) cells treated with LPS showed a strong activation of members of the MAPK family (pp38 and pERK) and their downstream substrates (pRSK, pCREB). While we also observed activation of pERK upon Poly(I:C) treatment, the peak response was significantly reduced (3.8 vs 16.8-fold increase above t=0) and delayed by 15 minutes. Activation of the mTOR complex is critical for proper microglial priming during inflammatory challenge (17). During our time-course we found that phosphorylated S6 kinase (pS6) levels showed a 25-fold increase after 1 hr of LPS-treatment. A similar pS6 upregulation was observed with Poly(I:C) but with delayed kinetics which likely point to differences in the early events following TLR4 or TLR3 engagement. Inflammatory markers such as pSTAT1 also showed a robust activation at 2 hrs with a similar degree of upregulation in both LPS and Poly(I:C) conditions. Persistent inflammatory challenge leads to alterations in microglial state (4). Indeed, with treatment of either LPS or Poly(I:C) we observed a progressive downregulation of Cx3CR1, a core microglia homeostatic marker. Towards the latter stages of the time-course we found upregulation of the microglial activation markers CD40, CD86, Ly6C, F480, and Galectin-1 with LPS producing more robust responses when compared with Poly(I:C). Notably, LPS, but not Poly(I:C), increased Ki67 expression, consistent with its known mitogenic effect on microglia (15) (Figure 1C, Supplementary Table 2A-B).

To corroborate some of our findings with more traditional methods for examining protein abundance, we performed western blots against pSTAT1/STAT1, pp38/p38, and pERK/ERK pairs for LPS-treated microglia. Overall, we found that our mass cytometry and immunoblot results corresponded well, further validating the use of mass cytometry to probe signaling pathways in microglial cells (Figure 2A-C, Supplementary Figure 2). Moreover, we found that our mass cytometry results demonstrated high reproducibility, as evidenced by strong agreement among experimental replicates (Figure 1B, Supplementary Figure 1B-C).

### High-dimensional analysis of microglial signaling profiles during inflammation

Microglia even in *in vitro* settings are heterogeneous. Our previous analysis examined the bulk CD11b^pos^CD45^pos^ response which provides a sense of general microglial signaling behavior. While useful, studying averaged responses does mask potentially interesting substructures that can only be elucidated at the single-cell level. To this aim, we performed leiden clustering on microglia using our panel and visualized the resulting clusters on a two-dimensional (2D) uniform manifold approximation and projection (UMAP) layout. We first clustered on cell identity markers and broadly categorized clusters corresponding to microglia and contaminating fibroblasts based on CD11b, CD45, GFAP, Olig2, and Fibronectin expression. A secondary round of clustering was then performed on the microglia using our signaling markers plus identity markers that may reflect activation state (Supplementary Figure 3A-B). Time-courses from both the microglia-only LPS and Poly(I:C) settings were clustered together. Following this second round of clustering, we identified 20 molecularly distinct clusters (Fig. 3A-D). We ordered clusters based on their relative abundance over the time course. For example, Cluster 1 cells were more abundant at early time-points and waned over the time-course while the opposite is true for cells in Cluster 20 (Figure 3D-F).

The predominant clusters at baseline exhibited a “signaling low” state with the exception of Cluster 3 which showed increased levels of the inflammatory and stress markers Galectin-1, pSrc, and pNFκB. Also noteworthy are Clusters 4 and 5 which had markedly lower Cx3CR1 levels than the other baseline clusters (Figure 3D, G). Previous work has demonstrated that microglia in culture exhibit significant heterogeneity and can down-regulate homeostatic markers such as *Cx3cr1* (18). Our observations are in-line with these activated subsets that can exist at baseline in cultured settings. Upon LPS stimulation, a transient pERK^hi^pp38^hi^pRSK^hi^pCREB^hi^ population (cluster 8) emerged at 15 minutes, comprising 73% of microglia before dissipating by 30 minutes. Poly(I:C) also induced this subset, but to a lesser extent and with delayed kinetics (Figure 3B-D, G). Consistent with our line-plots in Figure 1, we observed pSTAT1^hi^ clusters (Clusters 11 and 12) emerge at the 2 hr mark. However, considerably fewer LPS-treated cells seemed to pass through Clusters 11-12 in comparison to Poly(I:C) treatment (63 % vs 36 %, respectively). Moreover, we also found a larger persistence of the pSTAT1 signature in the Poly(I:C) treatment condition. These data suggest an enhanced pSTAT1 signature for Poly(I:C)-mediated microglial activation compared to LPS. By the 24-48 hr mark, a majority of the cells in both the LPS and Poly(I:C) groups fell under Cluster 20 which was characterized by high levels of CD40, CD86, pSrc, pNFκB, and pSTAT3 (Figure 3D-F). Overall, we find that both LPS and Poly(I:C) treatment induce a MAPK->pSTAT1->CD40/CD86 pathway with MAPK signaling playing perhaps a lesser role in Poly(I:C) activation.

### Presence of astrocytes dampens microglial pro-inflammatory signaling

Given that microglia do not exist in isolation when responding to challenges *in vivo*, we were interested in examining how other glial cells influence microglial signaling. To this aim, we treated mixed glial cultures (hereafter referred to as microglia + astrocytes) with LPS or Poly(I:C) and processed the samples for mass cytometry. Interestingly, when we performed these same experiments in the presence of astrocytes, we observed a striking blunting in microglial response across many markers (Figure 3B-C, Supplementary Figure 4A-B, Supplementary Figure 5A-B). Notably, CD40 which peaked with a 9-fold and 20-fold increase in our microglia cultures following LPS or Poly(I:C) addition was only increased by 1-2 fold in microglia + astrocyte cultures with either LPS or Poly(I:C) at the 24 hr time-point. Decreases were also seen with pS6, pCREB, pSTAT1, and pSTAT3 (Figure 4C).

We then performed high-dimensional analysis on all 4 of our datasets (Microglia LPS, Microglia Poly(I:C) (PIC), Microglia + Ast LPS, and Microglia + Ast PIC) first clustering the cells on identity markers then later performing secondary clustering on microglia as previously described (Supplementary Figure 3C-D). It is worth mentioning that the astrocytes in our culture were strongly adhered to the coated surface making their complete removal following accutase addition challenging. In order to best preserve cell signaling state we attempted to collect samples and place them in fixative in < 1 min. Because of this, the percentage of astrocytes collected likely underrepresent the actual percentage in culture (data not shown).

We observed 19 clusters which bore molecular similarity to the previously described 20 clusters in Figure 2. Overall, we did not observe significant differences between the apparent paths that microglia take when challenged with LPS or Poly(I:C) with the inclusion of astrocytes. In the presence of astrocytes, microglia treated with LPS and Poly(I:C) still proceed through a MAPK->pSTAT1->CD40/CD86 trajectory. However it is important to recognize that the expression of certain key markers show a potent down-regulation (e.g. CD40) with the presence of astrocytes resulting in a terminal state(s) that exhibit a decreased inflammatory profile (Supplementary Figure 6A). Many cells in all conditions reach Cluster 19 (CD40^hi^CD86^hi^pSTAT3^hi^pNFκB^hi^) by 24 hours, but significant divergence occurs at 48 hours. The percentage of cells in Cluster 19 were similar in LPS-treated microglia and microglia + astrocyte cultures (56.4 % and 53.4 %, respectively). However, Poly(I:C)-treated microglia + astrocytes had significantly fewer Cluster 19 cells when compared to Poly(I:C)-treated microglia (39.8 % vs 3.68 %) (Figure 4G-H). To gauge overall UMAP similarity at the 48 hr time-point, we computed the root-mean-square error (RMSE) in cluster abundance at the 0 and 48 hr marks for each of our conditions. Similar RMSEs were observed for LPS-treated microglia and microglia + astrocyte cultures while the RMSE from Poly(I:C)-treated microglia + astrocyte cultures was 3-fold lower. This suggests that microglia cultured in the presence of astrocytes following Poly(I:C) treatment display an increased capacity towards returning closer to baseline (Supplementary Figure 7A-D).

### Astrocyte-intrinsic signaling responses to inflammatory stimuli

To better understand the effects of astrocytes on microglial signaling, we evaluated astrocyte responses following LPS or Poly(I:C) challenge. We found that astrocytes exhibited a distinct signaling profile compared to microglia, with significantly higher levels of β-catenin, pAkt, pNFκB, pcJun, and pγH2AX across the duration of both LPS and Poly(I:C) treatments (Supplementary Figure 7A-B). Notably, β-catenin signaling, which has been shown to play a critical role in mediating astrocyte activation (19), was largely absent in microglia. Furthermore, we observed a rapid upregulation of pSTAT3 in astrocytes at the 1-hour time point, coinciding with the peak of microglial pSTAT1 activation. Interestingly, LPS induced a more robust pSTAT3 response in astrocytes compared to Poly(I:C) (Figure 5A-B), suggesting stimulus-specific regulation of this pathway.

The activation of STAT3 signaling in astrocytes is particularly relevant, as this pathway has been implicated in the induction of immunosuppressive factors such as IL-10 and TGF-β (20, 21). Moreover, reactive astrocyte subsets expressing high levels of STAT3 have been shown to exert neuroprotective effects in the context of CNS injury and inflammation (20, 22). Given the observation that microglia cultured in the presence of astrocytes exhibit dampened inflammatory responses, it is tempting to speculate that the astrocyte-derived factors downstream of STAT3 activation may contribute to the immunomodulation of microglia. However, further investigation, such as transcriptomic analysis of astrocytes or targeted perturbation of STAT3 signaling, is necessary to confirm this hypothesis and elucidate the precise mechanisms underlying the astrocyte-microglial crosstalk.

## Discussion

Microglia are uniquely positioned to sense and respond to perturbations in CNS homeostasis. Both acute and chronic neuroinflammatory conditions elicit profound changes in microglial state, many of which have been characterized at the transcriptional level (4, 6, 15, 16). However, a comprehensive understanding of microglial behavior requires a detailed examination of the signaling pathways that regulate their responses. In this study, we employed single-cell mass cytometry to interrogate the temporal dynamics of microglial signaling following stimulation with the bacterial endotoxin LPS and the viral mimetic Poly(I:C). By simultaneously monitoring 33 markers over a 48-hour time course, we uncovered distinct signaling trajectories and astrocyte-mediated immunomodulation of microglial activation.

Our data revealed that both LPS and Poly(I:C) induced rapid activation of MAPK signaling, followed by a delayed upregulation of pSTAT1. However, the magnitude of pp38, pERK, and pRSK activation was significantly lower in Poly(I:C)-treated microglia compared to LPS. These differences likely reflect the distinct signaling cascades downstream of TLR4 and TLR3, the respective receptors for LPS and Poly(I:C). TLR4 engages both MyD88 and TRIF adaptor proteins, with MyD88 recruiting IRAK4 and TRAF6 to activate TAK1 and drive MAPK and NF-κB signaling (14, 23, 24). In contrast, TLR3 signals exclusively through TRIF, which can activate TAK1 but also triggers IRF3-dependent responses (14, 24, 25). The diminished MAPK activation in Poly(I:C)-treated microglia suggests a preferential engagement of IRF3 over TAK1 downstream of TLR3, consistent with previous reports (24, 25). Furthermore, the delayed kinetics of MAPK activation following Poly(I:C) stimulation aligns with the slower activation of TAK1 by TRIF compared to MyD88 (24, 25). Both TLR3 and TLR4 can induce interferon production via TRIF, leading to autocrine/paracrine STAT1 activation (14, 25, 26), as observed in our experiments. Collectively, these findings highlight the power of mass cytometry to recapitulate known aspects of TLR signaling while providing a more nuanced view of pathway dynamics.

A hallmark of microglial activation in inflammatory and neurodegenerative disorders is the downregulation of homeostatic markers (e.g. Cx3cr1, P2ry12, Tmem119) and upregulation of molecules associated with pro-inflammatory and phagocytic functions (4). Consistent with this, we observed a progressive loss of Cx3CR1 expression and increased levels of the activation markers CD40 and CD86 in LPS- and Poly(I:C)-treated microglia. While both stimuli led to a CD40^hi^CD86^hi^ phenotype, only LPS induced a significant upregulation of F4/80 and Ly6C, suggesting stimulus-specific regulation of microglial states. These findings underscore the importance of examining microglial responses to diverse inflammatory triggers and caution against generalizing observations across stimuli.

Emerging evidence indicates that astrocytes play a critical role in modulating microglial responses in a context-dependent manner (22, 27–28). Intriguingly, we found that the presence of astrocytes markedly attenuated the inflammatory profile of LPS- and Poly(I:C)-stimulated microglia, with notable reductions in CD40, pS6, pCREB, pSTAT1, and pSTAT3 levels. This immunosuppressive effect was particularly striking for CD40, a key regulator of microglial activation in infectious and neurodegenerative diseases (29). Astrocyte-derived factors, such as IL-10 and TGF-β, have been shown to inhibit microglial *CD40* expression (28, 30), providing a potential mechanism for the observed immunomodulation. Importantly, despite the dampened inflammatory profile, microglia in mixed cultures still underwent MAPK and STAT1 activation, albeit to a lesser extent, and downregulated Cx3CR1, indicating a conserved core response to LPS and Poly(I:C). These findings suggest that astrocytes fine-tune the terminal activation state of microglia without fundamentally altering their signaling trajectories. Further investigation into the crosstalk between microglia and astrocytes, potentially through secretome profiling, could shed light on the molecular mediators underlying these effects.

How do astrocytes suppress microglial signaling response? While our data suggest that astrocyte-derived factors, such as IL-10 and TGF-β, may contribute to the suppression of microglial inflammatory responses, several other candidate pathways and mechanisms are worth considering. For instance, astrocytes have been shown to express high levels of the enzyme indoleamine 2,3-dioxygenase (IDO), which catalyzes the degradation of tryptophan into kynurenine (31). Kynurenine and its metabolites have potent immunosuppressive effects, and their production by astrocytes has been implicated in the regulation of microglial activation in the context of multiple sclerosis and Alzheimer’s disease (32, 33). Additionally, astrocytes are a major source of the anti-inflammatory mediator adenosine, which can attenuate microglial activation through the engagement of adenosine A2A receptors (34). Furthermore, astrocyte-derived exosomes have been found to contain microRNAs and other regulatory molecules that can modulate microglial function and suppress inflammation (35). Future studies employing transcriptomic and proteomic analyses of astrocyte-conditioned media, as well as targeted perturbation of these candidate pathways, will be essential in elucidating the precise mechanisms underlying the astrocyte-mediated immunomodulation of microglia.

While our study provides novel insights into microglial signaling dynamics, it is important to acknowledge its limitations. First, our findings are based on an in vitro model, which may not fully recapitulate the complex microenvironment of the CNS. Future studies using ex vivo or in vivo approaches, such as mass cytometry of acutely isolated microglia or imaging mass cytometry of brain tissue sections, could validate and extend our observations. Second, our CyTOF panel, while comprehensive, does not capture the entire spectrum of signaling pathways and microglial markers. Expanding the panel to include additional phospho-proteins (e.g., IRF3, TAK1), pro- and anti-inflammatory cytokines (e.g., TNFL, IL-10), and surface markers (e.g., P2ry12, Tmem119) could provide a more granular view of microglial states. Finally, our study focused on the effects of LPS and Poly(I:C), two commonly used inflammatory stimuli. Investigating microglial responses to other pathologically relevant triggers, such as amyloid-β, α-synuclein, or neuronal injury, could uncover stimulus-specific signaling patterns and potential therapeutic targets.

Our study demonstrates the power of single-cell mass cytometry to unravel the complex signaling landscape of microglia under inflammatory conditions. By providing a detailed view of the temporal dynamics and astrocyte-mediated regulation of microglial activation, our work lays the foundation for future mechanistic studies and highlights the potential of mass cytometry to identify novel therapeutic targets in neuroinflammatory disorders. As the field continues to embrace high-dimensional single-cell technologies, we anticipate that the integration of mass cytometry with transcriptomic, metabolomic, and imaging approaches will accelerate our understanding of microglial biology and unlock new avenues for therapeutic intervention.

## Data Availability

All datasets used and/or analyzed in this present study are available from the corresponding author upon request. The debarcoded sample FCS files are available at Cytobank (https://community.cytobank.org/cytobank/experiments/110288).

## Acknowledgements

We would like to thank all members of the Deppmann and Zunder labs for their helpful discussions. We would also like to specifically thank Corey Williams and Sarah Goggin of the Zunder lab for providing the code needed to run our analyses. Finally, we also thank the University of Virginia Flow Cytometry Core, RRID: SCR_017829 for technical assistance in antibody conjugation and with the CyTOF mass cytometer instrument.

## Author Contributions

SK and ADK were responsible for conceptualizing the project, designing experiments, and harvesting samples for mass cytometry. ABK performed all the antibody staining and submitted the samples for mass cytometry. SK performed the western blotting. SK and ADK performed all the data analysis. SK, CDD, and ERZ prepared figures and wrote the manuscript with input from all the authors.

## Figure Legends

**Supplementary Table 1**. Mass Cytometry Antibody Panel. Complete list of all markers used, concentrations of antibodies, and vendor information for mass cytometry studies.

**Supplementary Table 2**. P-values for Fig 1C comparing fold-change responses across time to time t=0 within a given condition (either LPS or Poly(I:C)). (A) Significant p-values following post-hoc corrections are provided for LPS and (B) Poly(I:C).

**Supplementary Figure 1.** Replicate analysis of microglia-only cultures treated with LPS or Poly(I:C). Gating strategy in Cytobank to isolate microglia. bc_separation_dist x mahalanobis_dist - gate to exclude non-barcoded events. Event_length x Center - gate to exclude doublet events as well as events that fall outside the normal Gaussian discrimination parameters for mass cytometry. 191Ir_Intercalator x Event_length - gate to remove non-cell events and dying/dead cells. 142Nd x 140Ce - gate to remove cerium contamination from normalization beads or environment. CD45 x CD11b - gate to select for microglia. CD45 x GFAP - secondary clean-up gate to remove residual astrocyte contaminants (more relevant for mixed cultures but was also applied to pure cultures to maintain consistency) (A). Histograms depicting marker counts for all pure replicates in LPS and Poly(I:C) conditions (B).

**Supplementary Figure 2.** Uncropped gels for Figure 2C.

**Supplementary Figure 3.** Isolating microglia for high-dimensional analysis. Cell events from microglia-only samples were exported from Cytobank and clustered on identity markers (A). Violin plot showing protein expression from clusters in (A). Clusters 7,11, and 15 were deemed to be non-microglia and excluded from secondary clustering (B).

**Supplementary Figure 4.** Summary of microglial LPS Responses. Fold-changes (A) or raw average intensities for all markers except CD11b, CD45, GFAP, and Olig2 (B) are shown for LPS-treated microglia-only and microglia+astrocyte cultures. Data were analyzed by two-way ANOVA followed by Šídák multiple comparisons test (A, B). Data are from at least three independent experiments and expressed as mean ± s.e.m. All data represent results taken from three independent cultures. Asterisks denote significance for Microglia vs Microglia + Ast responses at a given time-point. *p < 0.05, **p < 0.01, ***p < 0.001, ****p < 0.0001.

**Supplementary Figure 5.** Summary of microglial Poly(I:C) Responses. Fold-changes (A) or raw average intensities for all markers except CD11b, CD45, GFAP, and Olig2 (B) are shown for Poly(I:C)-treated microglia-only and microglia+astrocyte cultures. Data were analyzed by two-way ANOVA followed by Šídák multiple comparisons test (A, B). Data are from at least three independent experiments and expressed as mean ± s.e.m. All data represent results taken from three independent cultures. Asterisks denote significance for Microglia vs Microglia + Ast responses at a given time-point. *p < 0.05, **p < 0.01, ***p < 0.001, ****p < 0.0001.

**Supplementary Figure 6.** Comparison of Cluster 1 vs Cluster 19 marker intensity. Counts/cell for all markers used in secondary clustering are shown for cluster 1 and cluster 19 for all treatment conditions (A).

**Supplementary Figure 7.** Microglia from microglia + astrocyte Poly(I:C) setting show the greatest return to baseline after 48 hrs. Differences in cluster abundance between the 48 hr and 0 hr mark were calculated for microglia-only (LPS) (A), microglia + astrocyte (LPS) (B), microglia-only (Poly(I:C)) (C), and microglia + astrocyte (Poly(I:C)) conditions (D). Root-mean squared errors (using 0 as the expected value) are reported on lower-right.

