## Supplementary figures and images for "Characterizing microglial signaling dynamics during inflammation using single-cell mass cytometry"

### Supplemental Figure 1

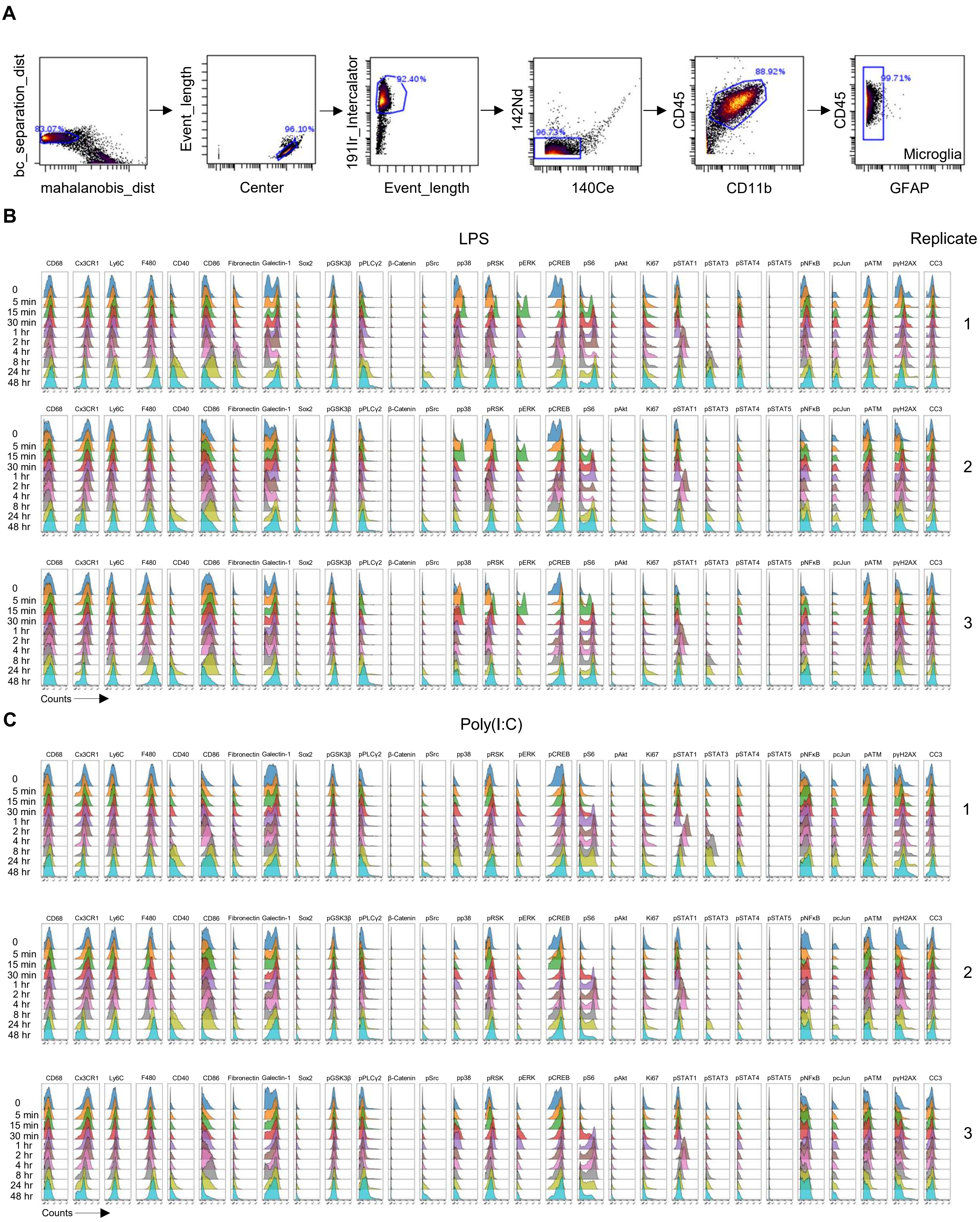

### Supplemental Figure 4

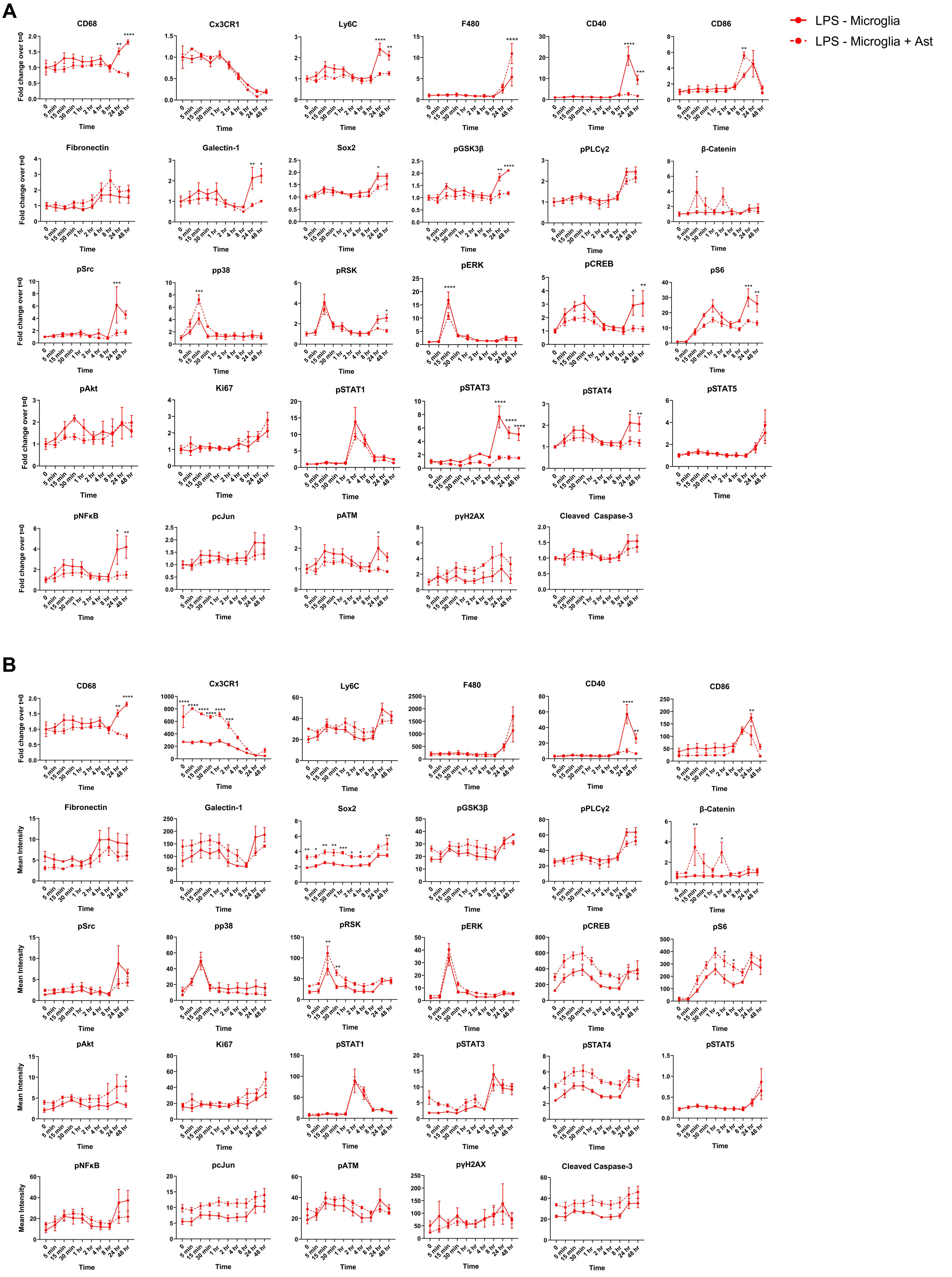

### Supplemental Figure 5

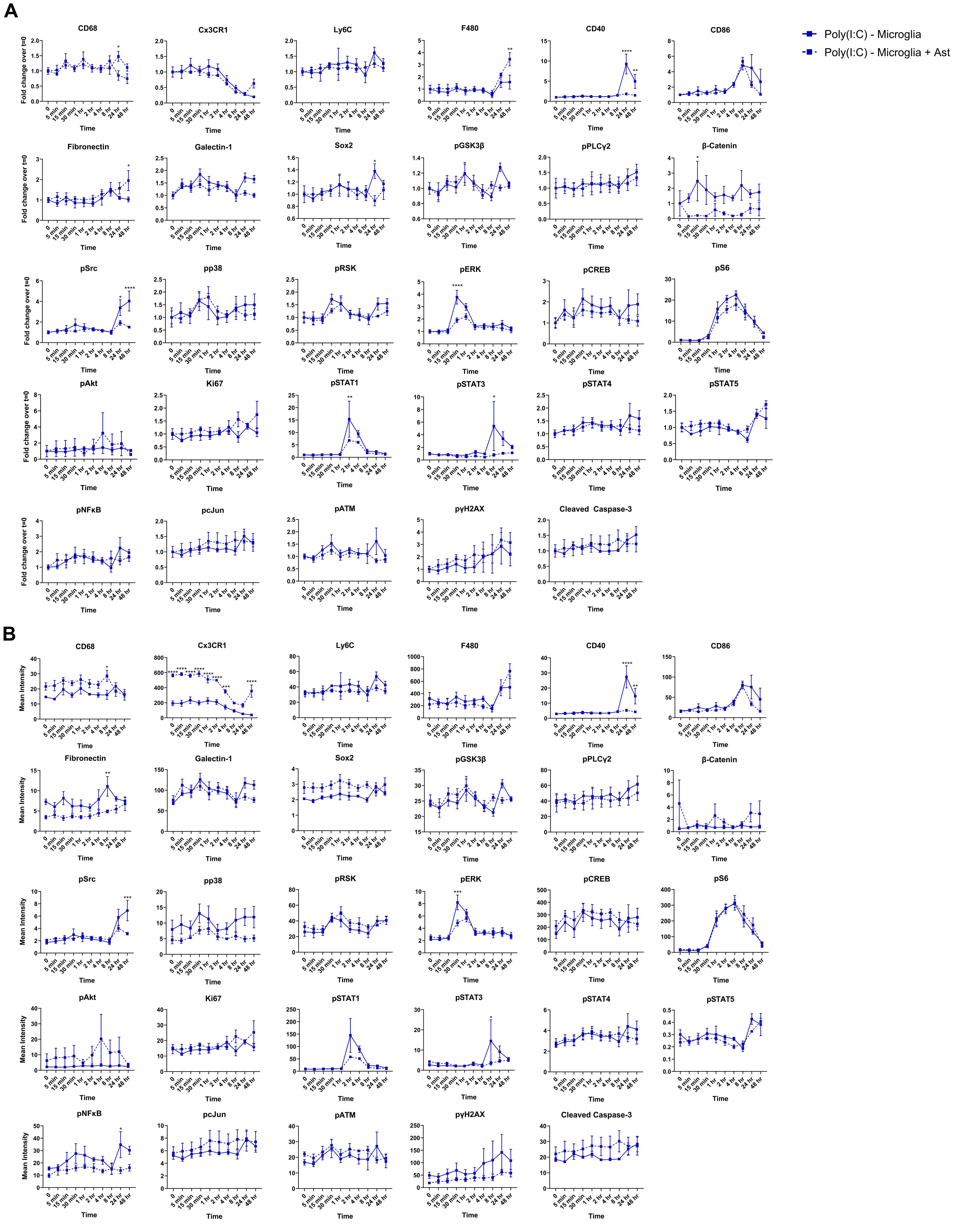

### Supplemental Figure 6

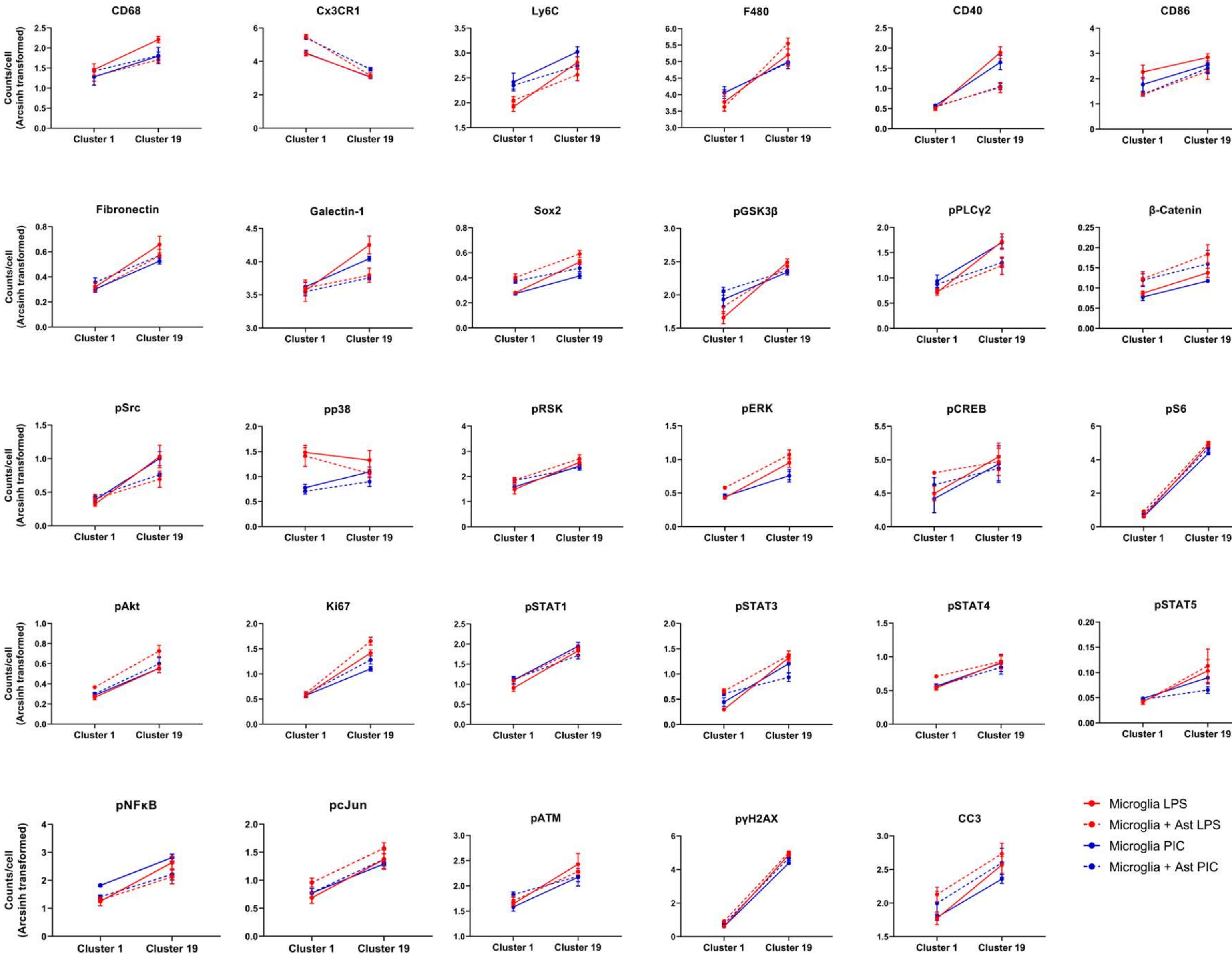

### Supplemental Figure 7

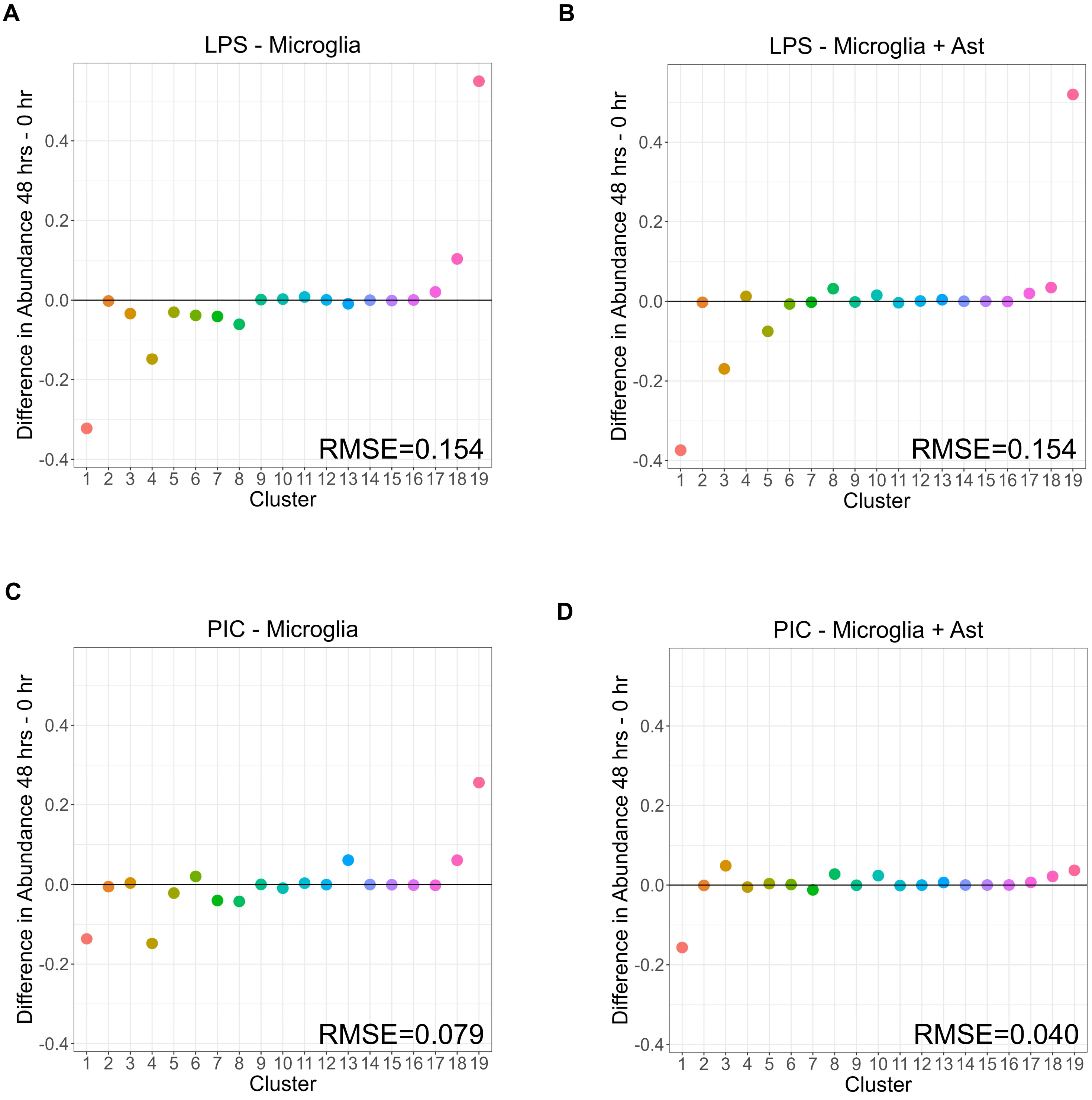
