## Supplemental Figure 2 for "Characterizing microglial signaling dynamics during inflammation using single-cell mass cytometry"

**Figure S2**

**A** Uncropped gels corresponding to Figure 2C

Order for all blots:

Ladder – Vehicle – 5 min LPS – 15 min LPS – 1 hr LPS – 2 hr LPS – 4 hr LPS – 24 hr LPS – Empty - Ladder

pp38

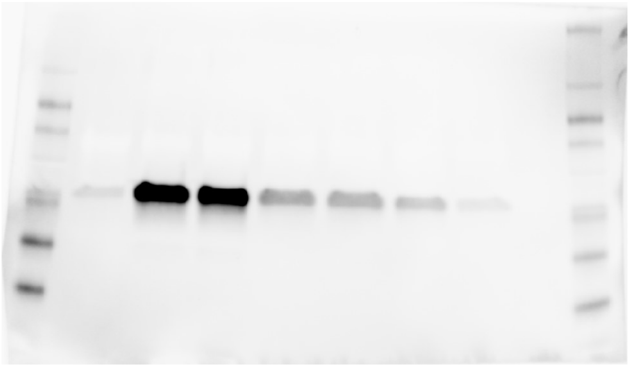

p38

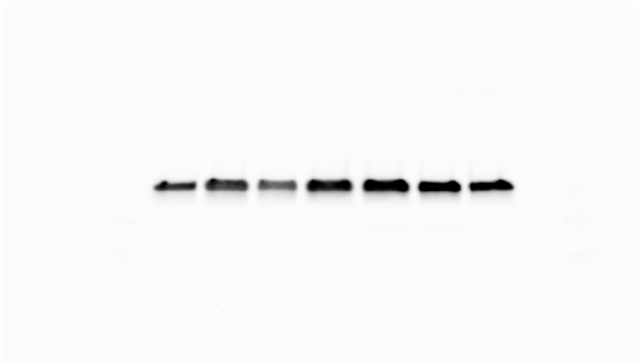

pERK

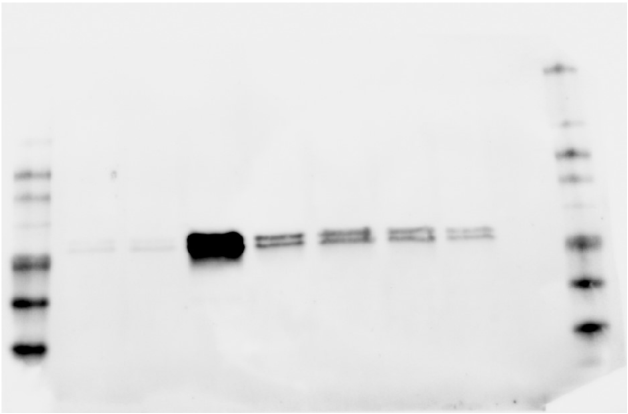

ERK

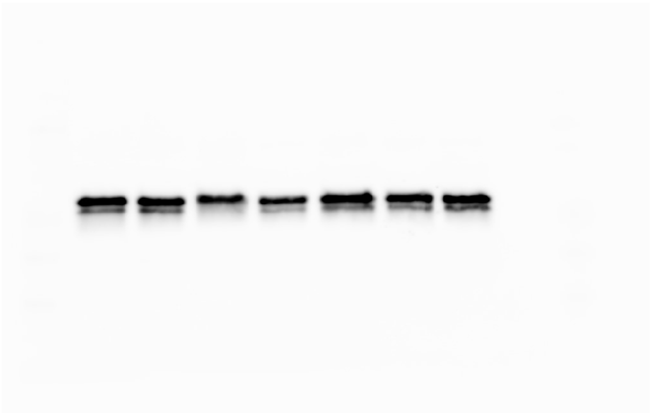

pSTAT1

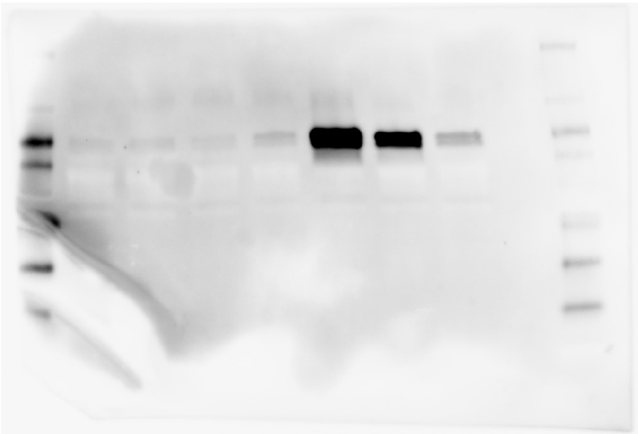

STAT1

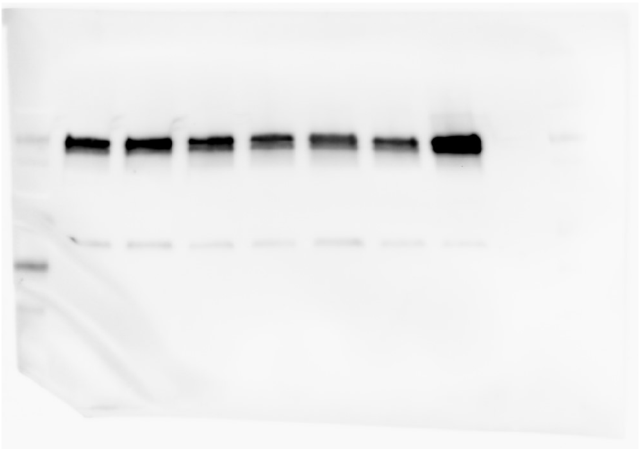
