## Supplemental Figure 3 for "Characterizing microglial signaling dynamics during inflammation using single-cell mass cytometry"

### Figure S3

**A**

1. Initial clustering on identity markers
2. Isolate microglia (CD11b$^{hi}$, CD45$^{hi}$, GFAP$^{lo}$, and Olig2$^{lo}$)
3. Recluster microglia on all markers except CD11b, CD45, GFAP, and Olig2

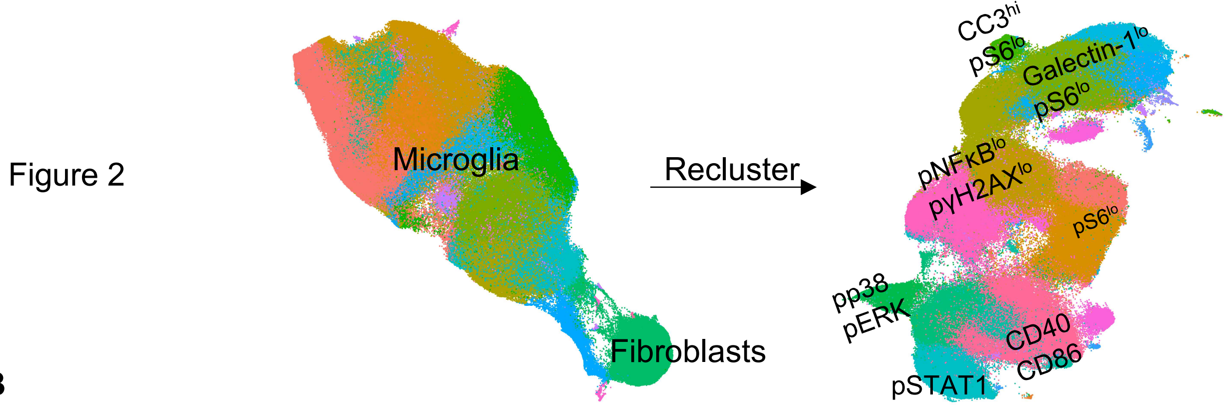


**B**

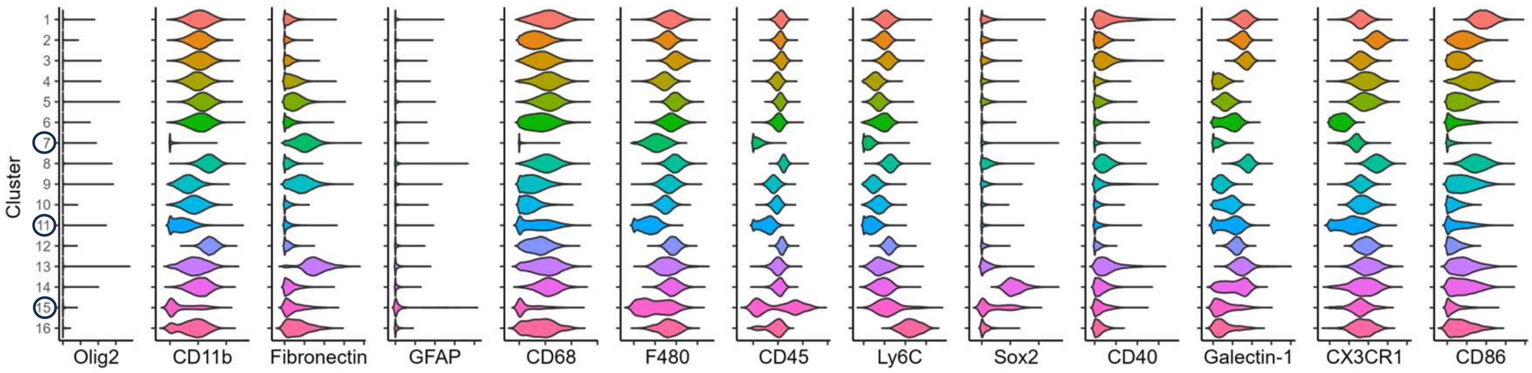


**C**

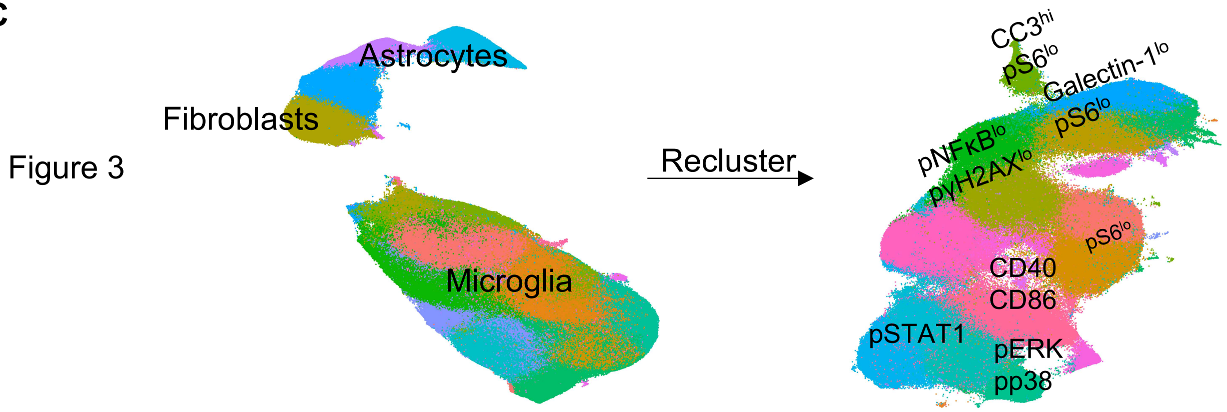


**D**

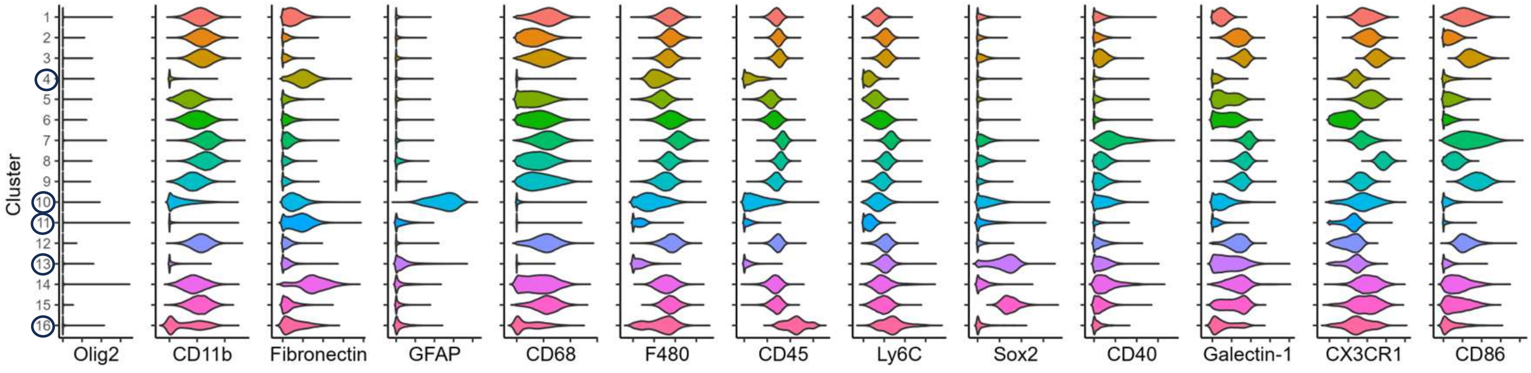


○ Denotes non-microglial clusters that were excluded for secondary round of clustering
