## Supplemental Table 1 for "Characterizing microglial signaling dynamics during inflammation using single-cell mass cytometry"

**Table S1**

**A**

### Mass Cytometry Antibody Panel

| Metal | Antigen | Concentration (ng/mL) | Vendor Information |
|---|---|---|---|
| In113 | Olig2 | 30 | Millipore (clone 211F1.1) |
| Cd114 | Cd11b | 55 | Biolegend (M1/70) |
| In115 | Fibronectin | 150 | BD Biosciences (610078) |
| La139 | Phospho-GSK3β (S9) | 1000 | CST (D85E12) |
| Pr141 | GFAP | 25 | BD Biosciences (556330) |
| Nd143 | CD68 | 25 | Biolegend (FA-11) |
| Nd144 | Phospho-PLCγ2 (Y759) | 2x | Fluidigm (K86-689.37) |
| Nd145 | Phospho-RSK (S320) | 2000 | R&D Systems (AF3369) |
| Nd146 | F4/80 | 1x | Fluidigm (BM8) |
| Sm147 | Phospho-STAT5 (Y694) | 1x | Fluidigm (47) |
| Sm149 | CD45 | 30 | Biolegend (30-F11) |
| Nd150 | Ly6C | 1x | Biolegend (HK1.4) |
| Sm152 | Ki67 | 2000 | Fluidigm (B56) |
| Eu153 | Phospho-STAT1 (Y701) | 1x | Fluidigm (58D6) |
| Sm154 | Phospho-Src (Y418) | 600 | BD Biosciences (K98-37) |
| Gd156 | Phospho-p38 (T180/Y182) | 1x | Fluidigm (D3F9) |
| Gd158 | Phospho-STAT3 (Y705) | 1x | Fluidigm (4/P-STAT3) |
| Tb159 | Phospho-ATM (S1981) | 700 | Biolegend (10H11.E12) |
| Gd160 | Sox2 | 1000 | R&D Systems (245610) |
| Dy161 | CD40 | 1x | Fluidigm (HM40-3) |
| Dy162 | Galectin-1 | 300 | R&D Systems (AF1245) |
| Dy164 | Cx3CR1 | 1x | Fluidigm (SA011F11) |
| Ho165 | Phospho-γH2AX (S139) | 700 | Biolegend (2F3) |
| Er166 | Phospho-NFκB (S529) | 1x | Fluidigm (K10x) |
| Er168 | Phospho-Akt (S473) | 2000 | BD Biosciences (560397) |
| Tm169 | Phospho-cJun (S73) | 5000 | CST (D47G9) |
| Er170 | βCatenin | 50 | CST (6B3) |
| Yb171 | Phospho-p44/42 MAPK (ERK1/2) (T202/Y204) | 1/3x | CST (D13.14.4E) |
| Yb172 | CD86 | 1x | Fluidigm (GL1) |
| Yb173 | CC3 | 5000 | BD Biosciences (C92-605) |
| Yb174 | Phospho-STAT4 (Y693) | 1/3x | Fluidigm |
| Lu175 | Phospho-S6 (S235/S236) | 500 | Fluidigm (N7-548) |
| Yb176 | Phospho-Creb (S133) | 1/3x | Fluidigm (87G3) |
