## Supplemental Table 2 for "Characterizing microglial signaling dynamics during inflammation using single-cell mass cytometry"

### Table S2

**A** **LPS**

| Marker | Šídák's multiple comparisons test | Adjusted p-value |
|---|---|---|
| CD68 | 0 vs 48 hr | 0.0019 (**) |
| Cx3CR1 | 0 vs 8 hr<br>0 vs 24 hr<br>0 vs 48 hr | 0.0050 (**)<br>0.0004 (***)<br>0.0002 (***) |
| Ly6C | 0 vs 24 hr<br>0 vs 48 hr | 0.0002 (***)<br>0.0051 (**) |
| F480 | 0 vs 48 hr | < 0.0001 (****) |
| CD40 | 0 vs 24 hr<br>0 vs 48 hr | < 0.0001 (****)<br>0.0005 (***) |
| CD86 | 0 vs 24 hr | 0.0031 (**) |
| Galectin-1 | 0 vs 24 hr<br>0 vs 48 hr | 0.0222 (*)<br>0.0085 (**) |
| Sox2 | 0 vs 24 hr<br>0 vs 48 hr | < 0.0001 (****)<br>< 0.0001 (****) |
| pGSK3β | 0 vs 15 min<br>0 vs 24 hr<br>0 vs 48 hr | 0.0239 (*)<br>< 0.0001 (****)<br>< 0.0001 (****) |
| pPLCγ2 | 0 vs 24 hr<br>0 vs 48 hr | < 0.0001 (****)<br>< 0.0001 (****) |
| pSrc | 0 vs 24 hr<br>0 vs 48 hr | 0.0002 (***)<br>0.0189 (*) |
| pp38 | 0 vs 15 min | 0.0002 (***) |
| pRSK | 0 vs 15 min<br>0 vs 24 hr<br>0 vs 48 hr | < 0.0001 (****)<br>0.0140 (*)<br>0.0044 (**) |

| Marker | Šídák's multiple comparisons test | Adjusted p-value |
|---|---|---|
| pERK | 0 vs 15 min | < 0.0001 (****) |
| pCREB | 0 vs 30 min<br>0 vs 48 hr | 0.0256 (*)<br>0.0298 (*) |
| pS6 | 0 vs 30 min<br>0 vs 1 hr<br>0 vs 2 hr<br>0 vs 4 hr<br>0 vs 8 hr<br>0 vs 24 hr<br>0 vs 48 hr | 0.0001 (***)<br>< 0.0001 (****)<br>0.0005 (***)<br>0.0199 (*)<br>0.0040 (**)<br>< 0.0001 (****)<br>< 0.0001 (****) |
| pAkt | 0 vs 30 min | 0.0095 (**) |
| Ki67 | 0 vs 48 hr | 0.0031 (**) |
| pSTAT1 | 0 vs 2 hr | 0.0004 (***) |
| pSTAT3 | 0 vs 8 hr<br>0 vs 24 hr | 0.0004 (***)<br>0.0448 (*) |
| pSTAT4 | 0 vs 24 hr<br>0 vs 48 hr | 0.0078 (**)<br>0.0142 (*) |
| pSTAT5 | 0 vs 48 hr | < 0.0001 (****) |
| pNFκB | 0 vs 24 hr<br>0 vs 48 hr | 0.0050 (**)<br>0.0020 (**) |
| pcJun | 0 vs 24 hr<br>0 vs 48 hr | 0.0200 (*)<br>0.0215 (*) |
| CC3 | 0 vs 24 hr<br>0 vs 48 hr | 0.0252 (*)<br>0.0207 (*) |

**B** **Poly(I:C)**

| Marker | Šídák's multiple comparisons test | Adjusted p-value |
|---|---|---|
| Cx3CR1 | 0 vs 8 hr<br>0 vs 24 hr<br>0 vs 48 hr | 0.0394 (*)<br>0.0012 (**)<br>0.0003 (***) |
| CD40 | 0 vs 24 hr | 0.0006 (***) |
| CD86 | 0 vs 8 hr<br>0 vs 24 hr | 0.0014 (**)<br>0.0039 (**) |
| pS6 | 0 vs 1 hr<br>0 vs 2 hr<br>0 vs 4 hr<br>0 vs 8 hr | 0.0015 (**)<br>< 0.0001 (****)<br>< 0.0001 (****)<br>0.0030 (**) |
| pSTAT1 | 0 vs 2 hr<br>0 vs 4 hr | < 0.0001 (****)<br>0.0365 (*) |
| pSTAT3 | 0 vs 8 hr | 0.0390 (*) |
| CC3 | 0 vs 48 hr | 0.0323 (*) |
